# Revealing S-layer Functions in the Hyperthermophilic Crenarchaeon *Sulfolobus islandicus*

**DOI:** 10.1101/444406

**Authors:** Changyi Zhang, Rebecca L. Wipfler, Yuan Li, Zhiyu Wang, Emily N. Hallett, Rachel J. Whitaker

## Abstract

The crystalline surface layer (S-layer), consisting of two glycoproteins SlaA and SlaB, is considered to be the exclusive component of the cell envelope outside of the cytoplasmic membrane in *Sulfolobus* species. Although biochemically and structurally characterized, the S-layer *in vivo* functions remain largely elusive in Archaea. Here, we investigate how the S-layer genes contribute to the S-layer architecture and affect cellular physiology in a crenarchaeal model, *Sulfolobus islandicus* M.16.4. Electron micrographs of mutant cells lacking *slaA* or both *slaA* and *slaB* confirm the absence of the outermost layer (SlaA), whereas cells with intact, partially, or completely detached SlaA are observed for the ∆*slaB* mutant. Importantly, we identify a novel S-layer-associated protein M164_1049, which does not functionally replace its homolog SlaB but likely assists SlaB to stabilize SlaA. Additionally, we find that mutants deficient in SlaA form large cell aggregates and the individual cell size varies significantly. The *slaB* gene deletion also causes noticeable cellular aggregation, but the size of those aggregates is smaller when compared to ∆*slaA* and ∆*slaAB* mutants. We further show the ∆*slaA* mutant cells exhibit more sensitivity to hyperosmotic stress but are not reduced to wild-type cell size. Finally, we demonstrate that the ∆*slaA* mutant contains aberrant chromosome copy numbers not seen in wild-type cells where the cell cycle is tightly regulated. Together these data suggest that the lack of *slaA* results in either cell fusion or irregularities in cell division. Our studies provide novel insights into the physiological and cellular functions of the S-layer in Archaea.

**Significance:** Rediscovery of the ancient evolutionary relationship between archaea and eukaryotes has revitalized interest in archaeal cell biology. Key to understanding the archaeal cell is the S-layer which is ubiquitous in Archaea but whose *in vivo* function is unknown. In this study, we genetically dissect how the two well-known S-layer genes as well as a newly identified S-layer-associated-protein-encoding gene contribute to the S-layer architecture in a hyperthermophilic crenarchaeal model *S. islandicus*. We provide genetic evidence for the first time showing that the *slaA* gene is a key cell morphology determinant and may play a role in *Sulfolobus* cell division or cell fusion.

## Introduction

The primary interface between the cell and its environment is a multi-functional cellular envelope. Comprised in this structure in some bacterial and most archaeal cells is a proteinaceous 2D-crystalline matrix coating the outside of the cell called the surface layer (S-layer). Despite its broad distribution in cells from two domains, a generalized function of the S-layer has not been identified. One unifying concept suggests the S-layer functions as an exoskeleton that interacts with other proteins in the cell membrane to coordinate diverse internal and external cell functions (1). In Bacteria, it has been shown that the S-layer is associated with either the peptidoglycan or the outer membrane (2), and S-layer genes are not essential for cellular viability in most studied bacteria under laboratory conditions. Characterization of S-layer mutants in Bacteria revealed that the S-layer plays highly diverse roles, serving as a protective coat or sieve, binding to specific receptors for adhesion, zones of adhesion for exoenzymes (1), maintaining cell envelope integrity (3), resisting osmotic stress (4), regulating cell morphology and contributing as a virulence factor (5), as well as maintaining cell swimming motility (6-8). In *Deinococcus radiodurans*, mutation in the S-layer protein encoding the gene *slpA* caused clumps of cells (3); however, these have not been investigated in detail. In contrast with the bacterial S-layers, archaeal S-layers are found to be the predominant if not the sole component of the cell wall outside of the cytoplasmic membrane with very few documented exceptions (9). So far, S-layer deletion mutants have not been successfully cultivated in any archaea until our recent report in a crenarchaeon *S. islandicus* (10), and archaeal S-layer studies has thus been limited to observational and biochemical analysis (9, 11) since it was discovered in a haloarchaea *Halobacterium salinarum* around 60 years ago (12).

Electron microscopy-based analyses of isolated proteinaceous S-layers in archaea revealed that they were organized as a highly regular two-dimensional lattice structure that displayed p2, p3, p4, and p6 symmetry depending on different species (9, 13). Moreover, it has been shown that the S-layer proteins in all studied archaea undergo post-translational modifications such as O-and N-glycosylation whereas the latter type is more prevalent (9, 11, 14). Apart from structural and biochemical studies, the *in vivo* functions of S-layer have not been studied extensively, but it has been proposed that the S-layer plays a role in osmotic stress (15), determines cell shape in a haloarchaeon *H. salinarium* (16), serves as a barrier to gene transfer in an isolated *S. islandicus* population (17), and contributes to cell stability as well as cell division in a methanogen *Methanocorpusculum sinese* (18).

Sulfolobales, within the Crenarchaeota whose cellular features have been suggested to have a close relationship to the last archaea-eukaryote common ancestor (19), is one of the predominant model systems where the S-layer structure is well understood. The structure of the S-layer in *Sulfolobus acidocaldarius* has been studied in great detail over the past thirty-five years. Electron microscopy analysis of the isolated cell wall sacculi in *S. acidocaldarius* revealed it was mainly composed of a single glycoprotein (known as the SlaA now), which was thought to exhibit a p6 symmetry (2) but was later confirmed to a p3 symmetry in this organism (20) and other studied *Sulfolobus* species (13). It is now well-known that the S-layer is composed of two glycosylated proteins, SlaA (~120 kDa) and SlaB (~45 kDa) in Sulfolobales (21-23). The current S-layer model in *Sulfolobus* shows a “stalk-and-cap” relationship between SlaA and SlaB, with SlaB as the stalk anchoring SlaA to the cytoplasmic membrane, forming a crystal matrix that constitutes the outermost layer covering the whole cell (22). Instability of the S-layer in *Sulfolobus* has been associated with changes in cell shape (24) and budding of vesicles (25, 26). It has been proposed that the archaeal S-layer assists the cell against turgor pressure (1, 9). Compensating for the absence of S-layer by forming a strong barrier at the site of cell division is hypothesized to be one role for Cdv proteins (27). The S-layer is believed to be a receptor for viruses and has been shown to change structure after viral induction and provide a barrier to virus egress during maturation of the SSV viral particle (28). No generalized function for the S-layer in Archaea has been defined as no archaeal S-layer deficient mutants have been characterized.

Recently, we have discovered that the S-layer genes were not essential for *S. islandicus* M.16.4 cell survival in standard lab conditions (10). Therefore, the resulting S-layer deletion mutants provide a model system to uncover the physiological and cellular roles of the archaeal S-layer. In this study, we aim at characterizing these S-layer deficient mutants to dissect *in vivo* functions of the S-layer in this model organism.

## Results

### Isolating roles for *slaA* and *slaB* in S-layer structure and function

Like other *Sulfolobus* species, in the genome of *S. islandicus* M.16.4, *slaB* is located in the downstream region of *slaA* with the same orientation (Supplementary Fig.1a). RT-PCR analysis showed that *slaA* and *slaB* are co-transcribed (Supplementary Fig. 1b), in agreement with a previous study in a related *Sulfolobus* species *S. solfataricus* P2 (22). To dissect how *slaA* and *slaB* contribute, separately and/or jointly, to the S-layer assembly in detail, we used electron microscopy to observe the three derivatives of *S. islandicus* RJW004 harboring an in-frame deletion of either *slaA* (∆*slaA*), *slaB* (∆*slaB*) or both genes (∆*slaAB*), as created recently (10). Conventional whole-cell transmission electron microscopy (TEM) analysis of the cell cultures revealed an intact outermost layer was retained around the whole cell in wild-type strain RJW004 (Fig. 1a and b). However, this outermost layer was not observed in any imaged ∆*slaA* (Fig. 1c and d) or ∆*slaAB* (Fig. 1g and h) mutant cells, indicating the SlaA contributes to the formation of this outermost layer (hereafter named SlaA). Intriguingly, a partial or discontinuous SlaA was frequently observed in the ∆*slaB* mutant cells (Fig. 1e and f), suggesting that SlaA was present on the outside of the cell but was fragile and unstable, and thus was easily detached from the cytoplasmic membrane following TEM sample preparation without the support provided by the SlaB stalk. These observations are in agreement with those obtained from our thin-section TEM experiment, as described previously (10). Interestingly, it is clear that cell surface appendages are expressed with visible pili and archaellum in all mutants (Fig. 1 and Supplementary Fig. 2).

**Figure 1.**
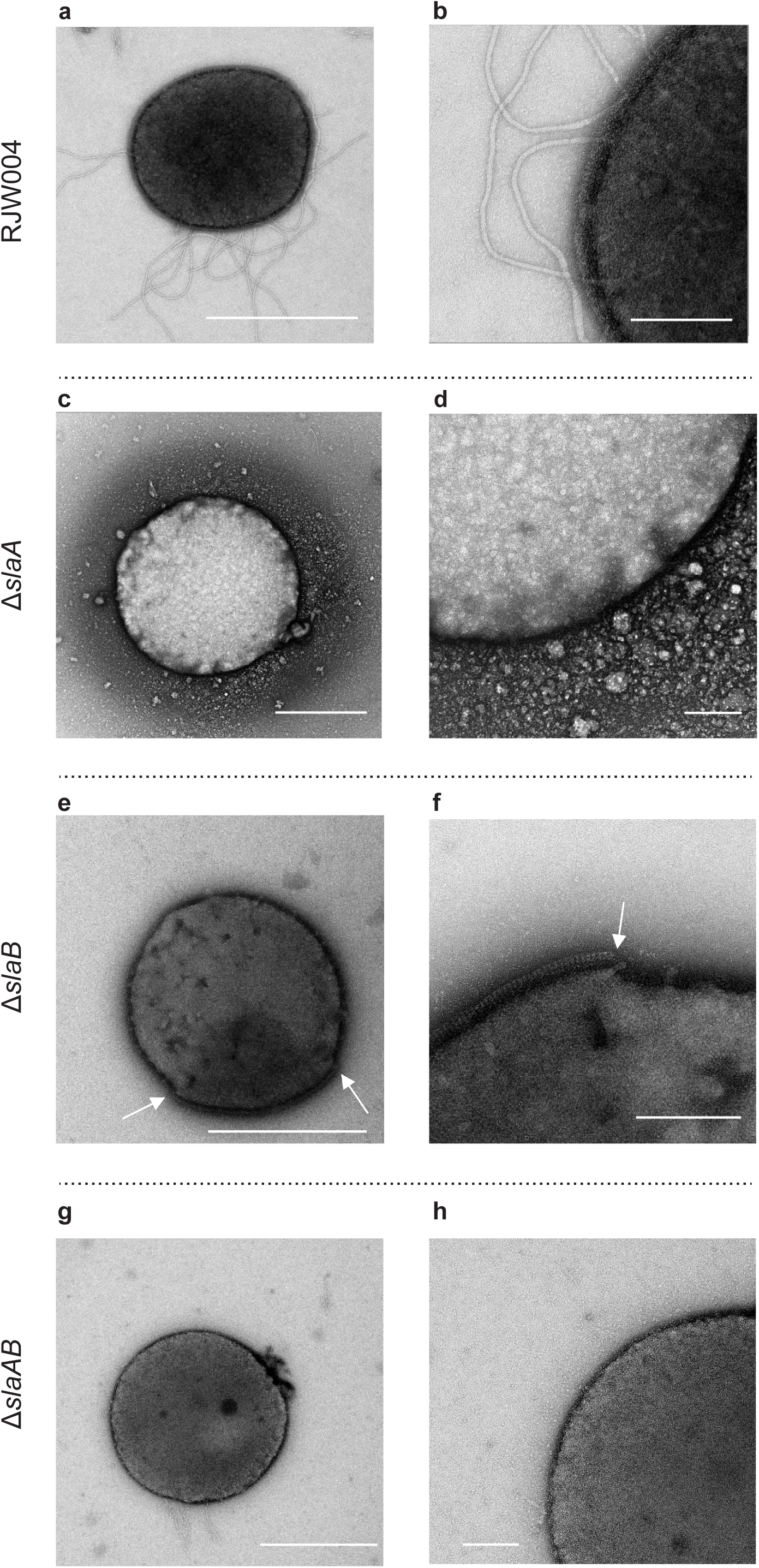
Conventional TEM analysis of the wild-type RJW004 and S-layer gene knockout mutants. RJW004 (**a-b**), △*slaA* (**c-d**), △*slaB* (**e-f**), △*slaAB* (**g-h**). **b** is a close-up image of **a**, **d** is a close-up image of **c**, **f** is a close-up image **e**, **h** is a close-up image of **g**. Breaking points of the outermost layer SlaA are indicated with white arrowheads. Scale bars, 1 μm (**a**, **c**, **e**, and **g**) and 200 nm (**b**, **d**, **f**, **h**, and **j**).

Next, we further examined the whole-cell morphology of S-layer mutant strains using scanning electron microscopy (SEM). As shown in Fig. 2a, 2b and Supplementary Fig. 3a, in all imaged cells of the wild-type strain RJW004, a regular lattice structure, characteristic of the S-layer, completely covers the whole cell surface. In contrast, this outermost lattice layer was not observed in the ∆*slaA* (Fig. 2c-e and Supplementary Fig. 3b) and ∆*slaAB* mutant cells (Fig. 2i-l and Supplementary Fig. 3d), further experimentally proving that SlaA makes up the sole component of the matrix layer that surrounds the cytoplasmic membrane. It was worth mentioning that no SlaA homologs are found when searching the genome of *S. islandicus* M.16.4, excluding the possibility that other protein could functionally complement SlaA. For the ∆*slaB* mutant strain, the outermost layer appeared to be peeled off partially (Fig. 2f and Supplementary Fig. 3c) or completely (Fig. 2g and Supplementary Fig. 3c) from the cell surface in most situations; however, cells with an intact outermost layer also existed (Fig. 2h), though observed rarely. These data further validate the idea that SlaB functions as a stalk to stabilize and tether SlaA to the cell membrane but suggest other proteins may serve this function as well. Noticeably, the cell surface composition appeared to be similar in both ∆*slaA* and ∆*slaAB* mutants, which was also indistinguishable from that of ∆*slaB* mutant cells in places where the SlaA protein layer was not attached (Fig. 2g).

**Figure 2.**
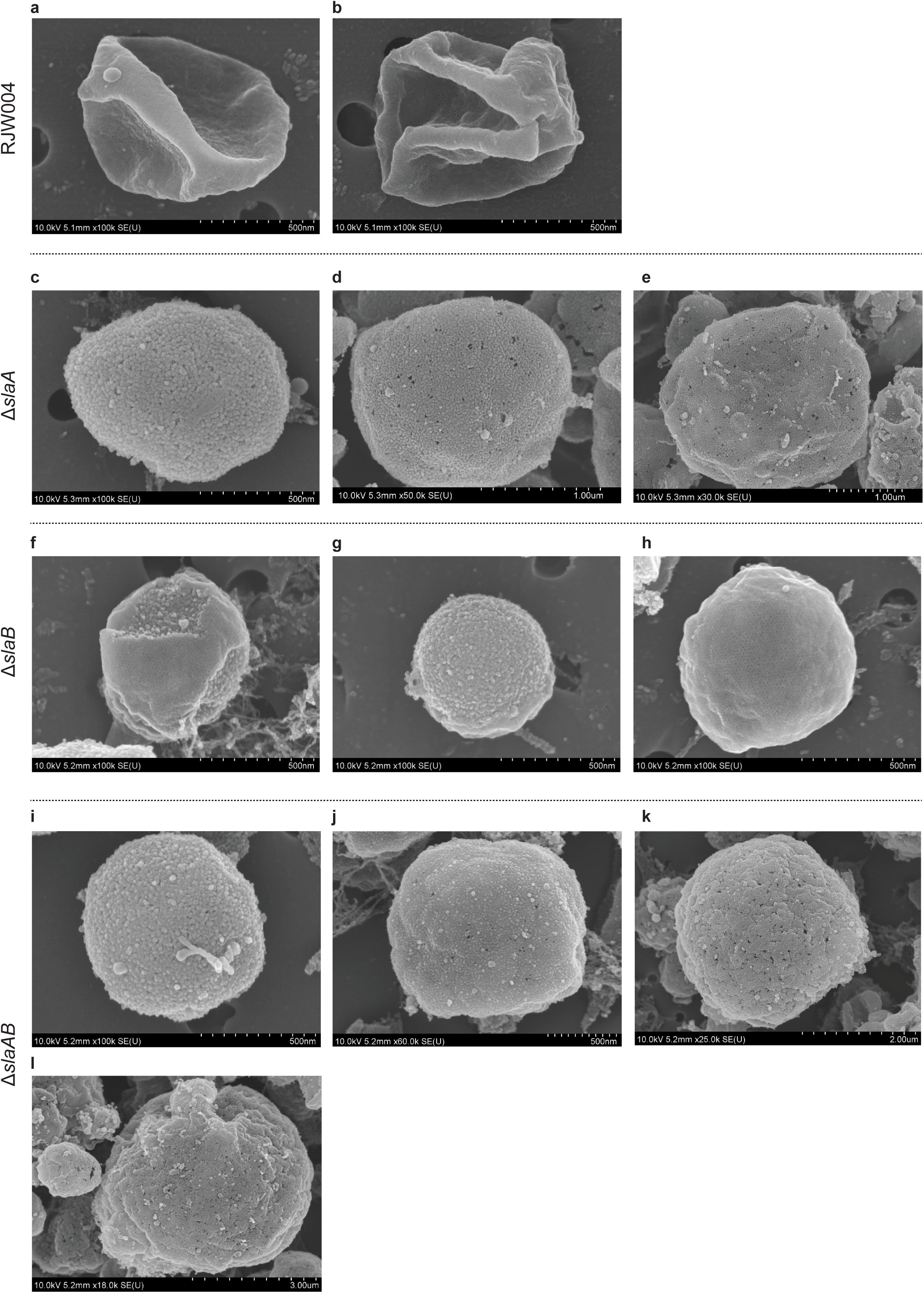
SEM analysis of wild-type RJW004 and S-layer gene knockout strains. Scanning electron micrographs showing the wild-type cell (**a-b**), ∆*sla*A mutant cell with no outermost layer and distinct sizes (**c-e**), ∆*slaB* mutant cell with partial (**f**), no (**g**), and intact (**h**) outermost layer, and ∆*slaAB* mutant cell with no S-layer and distinct sizes (**i-l**). Scale bars and image magnifications are shown in each micrograph. Additional SEM images are provided in the Supplemental Fig.3.

The cell size in SlaA^-^ mutants (∆*slaA* and ∆*slaAB*) varied significantly (Fig. 2c-2e, 2i-2l, and Supplementary Fig. 3b and 3d), ranging from ~0.7-5.0 μm, and the SlaA^-^ cells showed a morphology that was consistently unfolded to look markedly different from that of wild-type cells that have an irregular coccoid morphology (Supplementary Fig. 3a) as typically observed for *Sulfolobus* species (29, 30). The ∆*slaB* mutant cells varied slightly in size (~0.6-0.8 μm; Fig. 2f-2h) but showed an unfolded morphology (Supplementary Fig. 3c) consistent with the firm anchoring of the SlaA protein layer in the membrane being necessary for the rigidity that causes the irregular shape after fixation. This observation was also seen in thin-section TEM micrographs in our previous study (10) as well as in *S. solfataricus* with a structurally unsound cell envelope under high pressure freezing thin section TEM (24).

### M164_1049, a homolog of SlaB, does not functionally replace SlaB, but likely assists SlaB to stabilize the outermost layer SlaA

All electron microscopy (TEM, thin-section TEM, and SEM) analyses, presented both here and in a previous study (10), revealed a partial outermost layer was retained around the cell in the ∆*slaB* mutant, raising a question of whether there are homologs of SlaB that could fulfill the SlaB function. Using the SlaB protein sequence as a query, we identified a protein, M164_1049, annotated as a hypothetical protein in the genome, which shared 53% of amino acid identity with SlaB with 42% query coverage (Supplementary Fig. 4). We next investigated whether M164_1049 plays a role in S-layer retention, and particularly asked whether M164_1049 could partially or fully complement SlaB. To this end, we introduced a ∆*M164_1049* allele into the wild-type RJW004 and ∆*slaB* mutant strains using a recently developed microhomology-mediated gene inactivation strategy (31), which resulted in the single ∆*M164_1049* and double ∆*slaB*∆*M164_1049* mutants, respectively. The *M164_1049* deletion in mutant strains was confirmed by PCR analyses with the flanking and internal primer sets (Fig. 3a and 3b). The growth profile of ∆*M164_1049* mutant has no difference at all from its parental strain RWJ004 in that they exhibited the same growth rate and could reach approximately the same terminal OD_600nm_ (Fig. 3c). The double deletion mutant, ∆*slaB*∆*M164_1049*, displayed a comparable growth rate to its parental strain ∆*slaB* (Fig. 3d).

**Figure 3.**
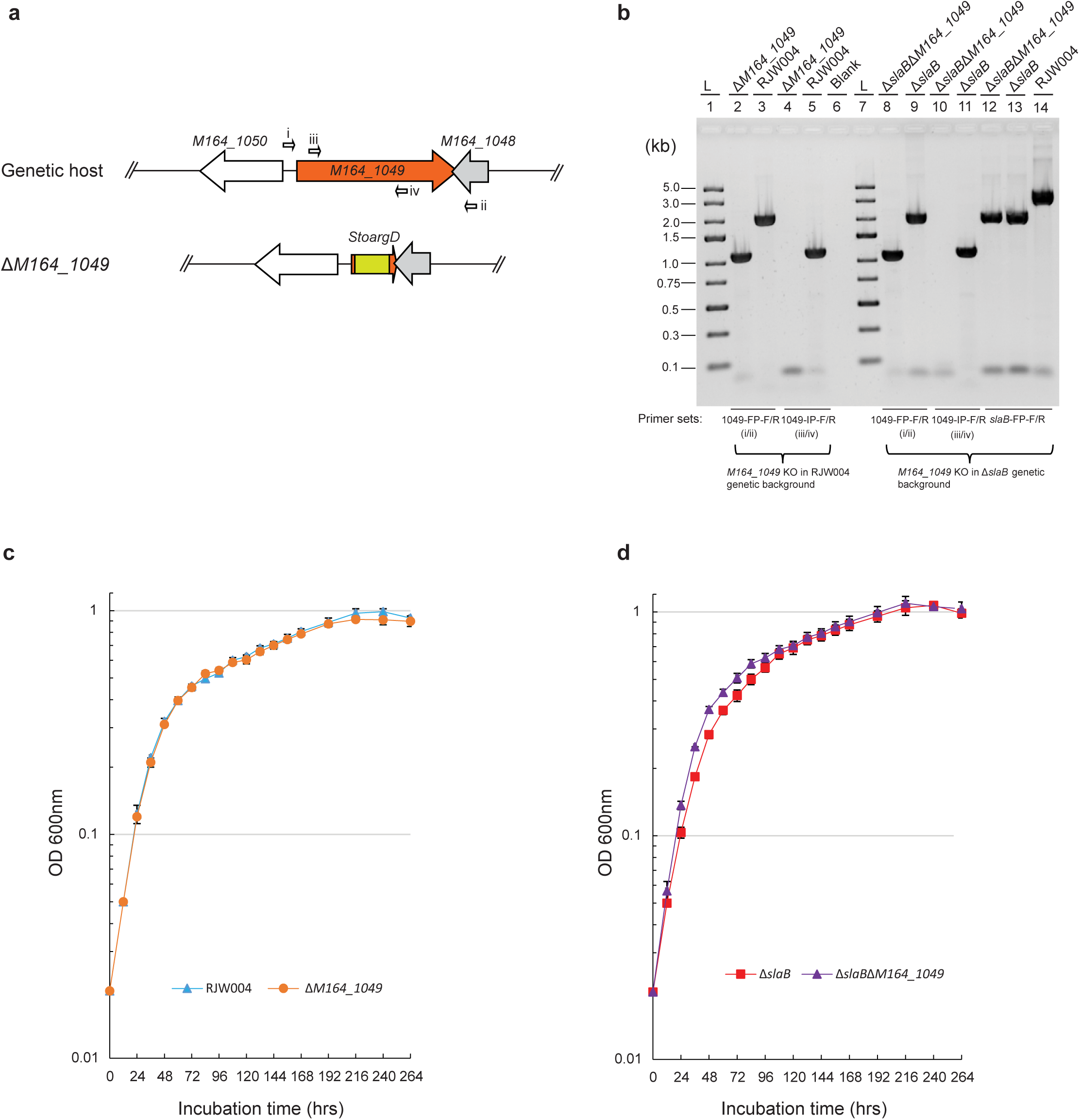
Characterization of *S. islandicus* mutants lacking the *M164_1049*. **a.** Genetic context of *M164_1049* in the genome of genetic host and mutant strain. The *M164_1049* in two host strains RJW004 and ∆*slaB* is replaced by an agmatine selection marker *StoargD* via homologous recombination as described previously(31). **b.** Confirming the deletion of *M164_1069* in host strains RJW004 and ∆*slaB*. *M164_1049* deletion is validated by PCR diagnoses with flanking primers binding outside the region of homology, and internal primers binding inside the region of *M164_1049*. The ∆*slaB* mutant allele is further confirmed in the ∆*slaB*∆*M164_1049* double deletion mutant using the *slaB* flanking primer set *slaB*-FP-F/R (lane 12). Relative positions of primers are indicated with small arrows. L (lanes 1 and 6) indicates the Quick-Load® 1 kb DNA Ladder (NEB, USA), with the size corresponding to each band labeled. Blank (lane 6) indicates no sample is loaded in the well when running the agarose gel electrophoresis. **c-d.** Growth curves of the ∆*M164_1049* deletion mutants and their parental strains. Strains are cultivated aerobically at 78 °C without shaking, and the cell optical density at 600 nm (OD_600nm_) are measured every 12 or 24 hours. Triple replicates are set up for each strain. Error bars represent SD.

To investigate whether M164_1049 contributes to the maintenance of S-layer, we used electron microscopy to observe the ∆*M164_1049* and ∆*slaB*∆*M164_1049* mutant cells. As shown in the whole-cell TEM analysis of the ∆*M164_1049* mutant, an intact and smooth outermost layer i.e. SlaA was consistently observed to encompass the whole cell (Fig. 4a and 4b), which was further validated by both thin-section TEM (Fig. 4c and 4d) and SEM (Fig. 4i, 4j, and Supplementary Fig. 5a) analyses. These observations were very similar to that of wild-type strain RJW004, indicating that M164_1049 does not contribute as a structural component of the S-layer lattice and should not function identically to SlaB. In contrast to the ∆*slaB* mutant cells in which a partial coat of SlaA was frequently observed, the SlaA of the ∆*slaB*∆*M164_1049* mutant was not observed surrounding the cells in all electron microscopy analyses (Fig. 4e-h and 4k-m). Interestingly, an increased amount of debris was observed in the SEM experiment for this double deletion mutant (Fig. 4k-m, and Supplementary Fig. 5b), which might correspond to unbound SlaA and other extracellular components. These cells, lacking both SlaB and M164_1049, may have been very sensitive to physical breaks/damages occurring in the process of SEM sample preparation. Even so, these results suggest that M164_1049 is indispensable to assist SlaB to firmly anchor the SlaA to the cytoplasmic membrane in *S. islandicus*.

**Figure 4.**
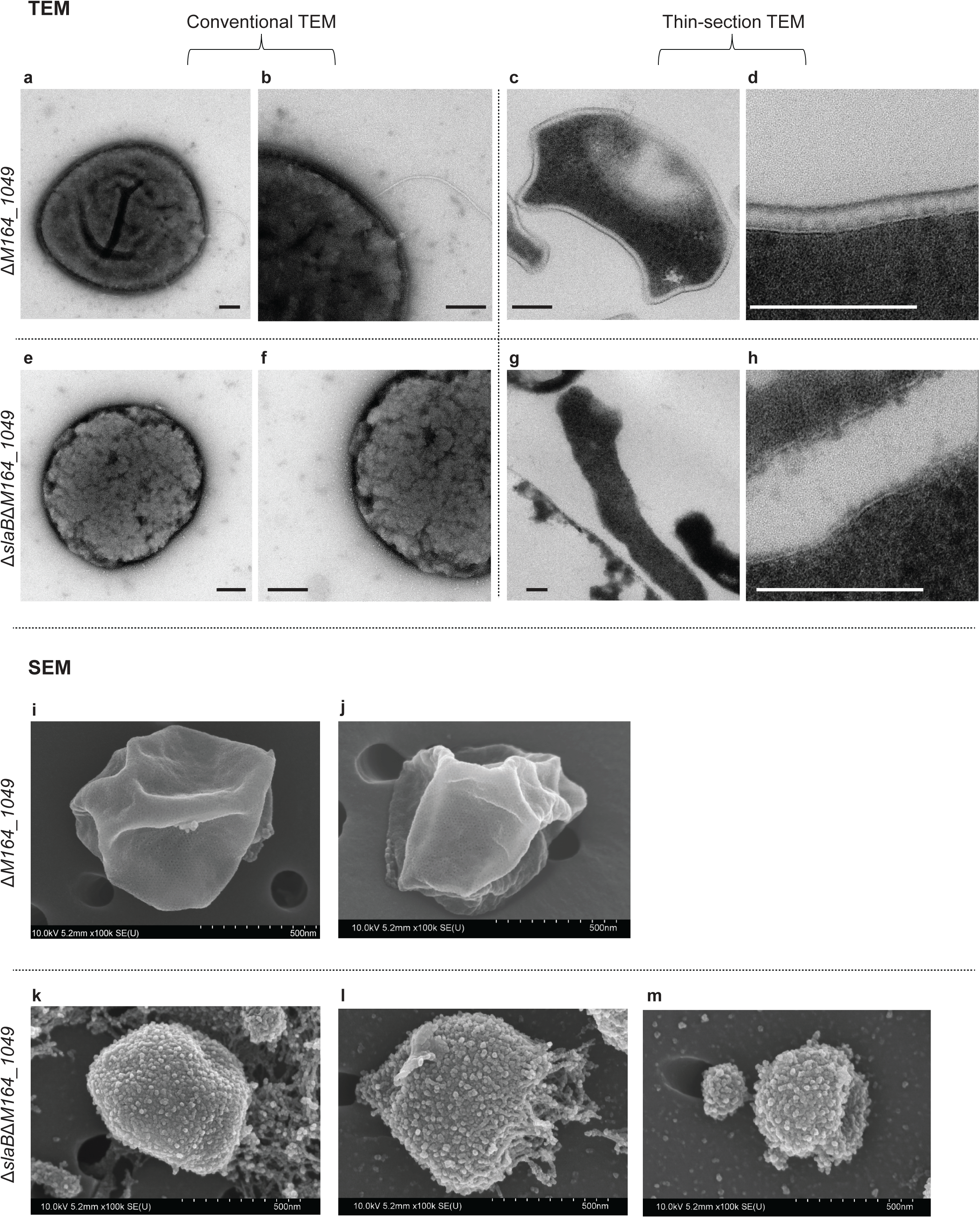
Electron micrographs of the ∆*M164_1049* and ∆*slaB*∆*M164_1049* mutant cells. Conventional TEM analysis of the ∆*M164_1049* (**a-b**) and ∆*slaB*∆*M164_1049* (**e-f**) mutants. Thin-section TEM analysis of the ∆*M164_1049* (**c-d**) and ∆*slaB*∆*M164_1049* (**g-h**) mutants. Images **b**, **f**, **d**, and **h** are close-up images of **a**, **e**, **c**, and **g**, respectively. Scale bars are 200 nm. Scanning electron micrographs of ∆*M164_1049* (**i-j**) and ∆*slaB*∆*M164_1049* (**k-m**) mutants. Scale bars and image magnifications are shown in each micrograph. Additional SEM images can be found in the Supplemental Fig. 5.

### Deficiency in SlaA results in large aggregates with variation in cell size

To characterize the phenotypes of these S-layer deficient mutants, cell cultures taken from log phase were observed under a light microscope. Surprisingly, compared to the wild type (Fig. 5a), we found that the cells predominantly form bulky clumps in the ∆*slaA* mutant cultures, generally containing hundreds or thousands of cells in a single clump or aggregate (Fig. 5b). Notably, we observed that the cell size within the aggregates varied significantly in the ∆*slaA* mutant, ranging from ~1-9 μm (Supplementary Fig. 6), which is normally larger than that observed in SEM (~0.7-5.0 μm). Deletion of the single *slaB* gene caused small aggregates in the cell cultures (less than 100 cells per aggregate), and the cell size within the aggregates varied, though not as much as the ∆*slaA* mutant, with a range of ~0.8-3.5 μm (Supplementary Fig. 7). Again, the observed cell size of the ∆*slaB* mutant under light microscopy is larger than that observed in SEM (~0.6-0.8 μm).

**Figure 5.**
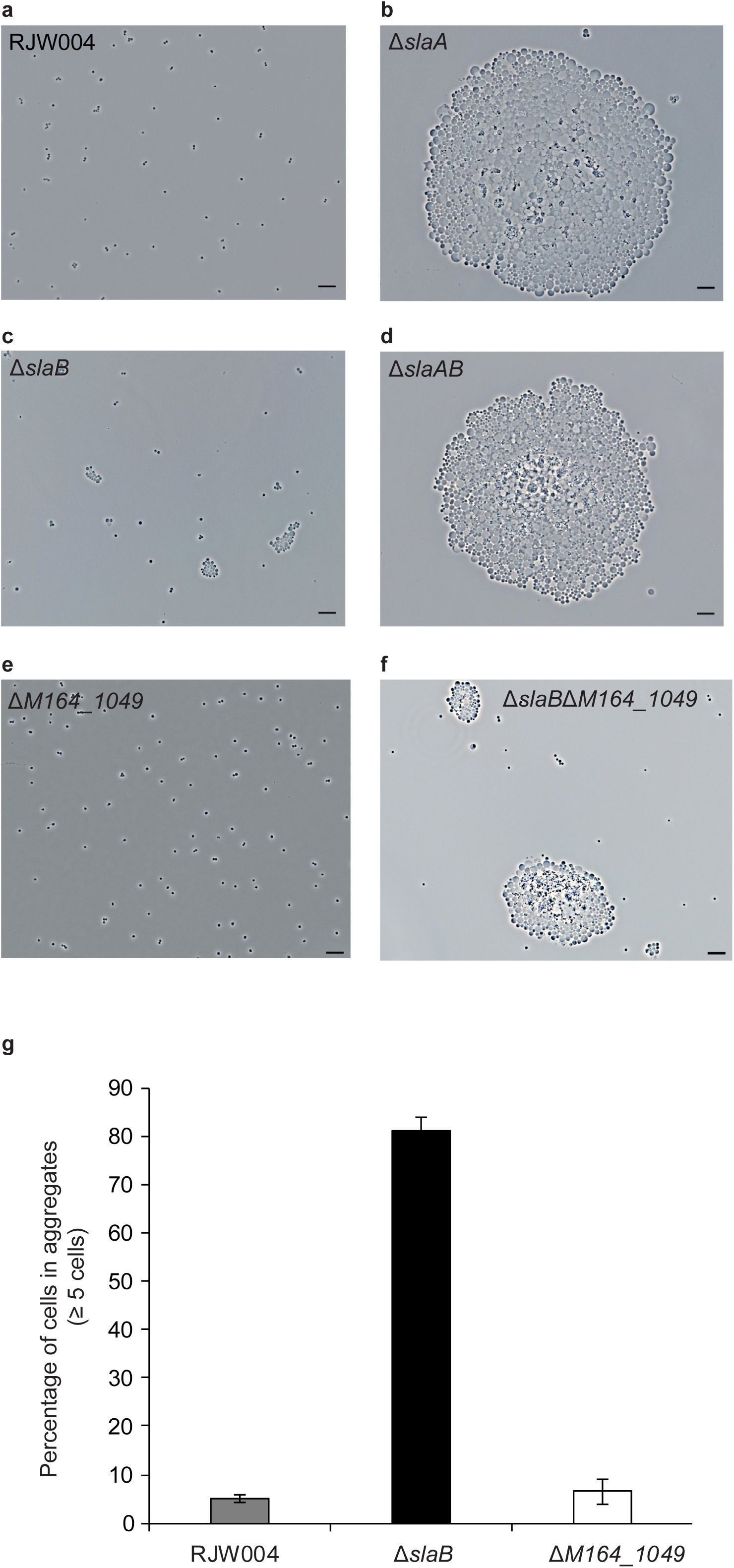
Phenotypic characterization of S-layer gene knockout mutants under a phase contrast microscope. **(a-f)** Representative phase-contrast microscopy images of cells from wild-type RJW004, ∆*slaA*, ∆*slaB*, ∆*slaAB*, ∆*M164_1049*, and ∆*slaB*∆*M164_1049* mutants, respectively. Five microliter of cell cultures were spotted on a cleaned microscope slide, covered with a coverslip, and then observed under a microscope. Scale bars, 10 μm. **(g)** Quantification of levels of cellular aggregation in wild-type RJW004, ∆*slaB*, and ∆*M164_1049* mutant cell cultures. At least 1,000 individual and aggregated cells from each sample were manually counted. Three biological replicates for each sample were conducted. Error bars represent SD.

Like the ∆*slaA* mutant, a substantial number of large clumps containing large cells was also observed in the ∆*slaAB* mutant cultures (Fig. 5d). The phenotypes, such as cell size and shape, between ∆*slaA* and ∆*slaAB* mutants are hardly distinguishable; however, the size of clumps in the ∆*slaAB* mutant is generally smaller than that of the ∆*slaA* mutant alone suggesting a role for *slaB* in cell aggregation. No obvious cellular aggregation was seen in the single ∆*M164_1049* mutant cells (Fig. 5e), similar to that in wild-type strain RJW004 (Fig. 5a). Interestingly, the size of the aggregate increased in the ∆*slaB*∆*M164_1049* double mutant (Fig. 5f) compared to that of the ∆*slaB* mutant. Large cellular clumps in the ∆*slaB*∆*M164_1049* mutant at a similar scale as in the ∆*slaA* and ∆*slaAB* mutants were occasionally observed (data not shown).

Further quantification of the level of cellular aggregation in the ∆*slaB* mutant cultures showed that approximately 80% of cells are in the aggregates (≥5 cells/aggregate) whereas only ~5% and ~6.5% of cells form aggregates (≥5 cells/aggregate) in the wild-type RJW004 and ∆*M164_1409* mutant cultures (Fig. 5g), respectively. Precisely evaluating the extent of cellular aggregation was challenging for the ∆*slaA*, ∆*slaAB*, and ∆*slaB*∆*M164_1409* mutants due to the difficulty in counting the exact number of cells in the larger clumps.

### *S. islandicus* cells deficient in S-layer still maintain cell membrane integrity

As addressed above, *Sulfolobus* cells deficient in the major S-layer component SlaA, either with the *slaA* gene deleted from the chromosome (∆*slaA* or ∆*slaAB* mutant) or having the wild-type *slaA* allele but no SlaA layer observed (∆*slaB*∆*M164_1409* mutant), showed a significant change in cell morphology. Thus, in following studies, we focused on the investigation of the single ∆*slaA* deletion mutant to determine the roles of *slaA* concerning cellular morphology and function. First, we evaluated the cell viability of the ∆*slaA* mutant by spotting assays, and it showed that the ∆*slaA* mutant exhibits more than 10-fold lower cell viability than that of wild-type strain (Fig. 6a). Next, we used live-dead fluorescence microscopy to see whether the aggregated cells in large clumps are alive or dead, and it was revealed most of the aggregated cells in the ∆*slaA* mutant are not permeable by propidium iodide (Fig. 6b). This indicates that they still maintained an impermeable cell membrane, suggesting the majority of the cells are alive. The large, irregular red shapes in the aggregates are supposed to be the lysed cells whose membrane was disrupted when undergoing imaging, a phenomenon not observed in wild-type cell (Fig. 6c). Interestingly, many of the larger cells had a decrease in staining in the center of the cells (Supplementary Fig. 8 and Supplementary Movie 1).

**Figure 6.**
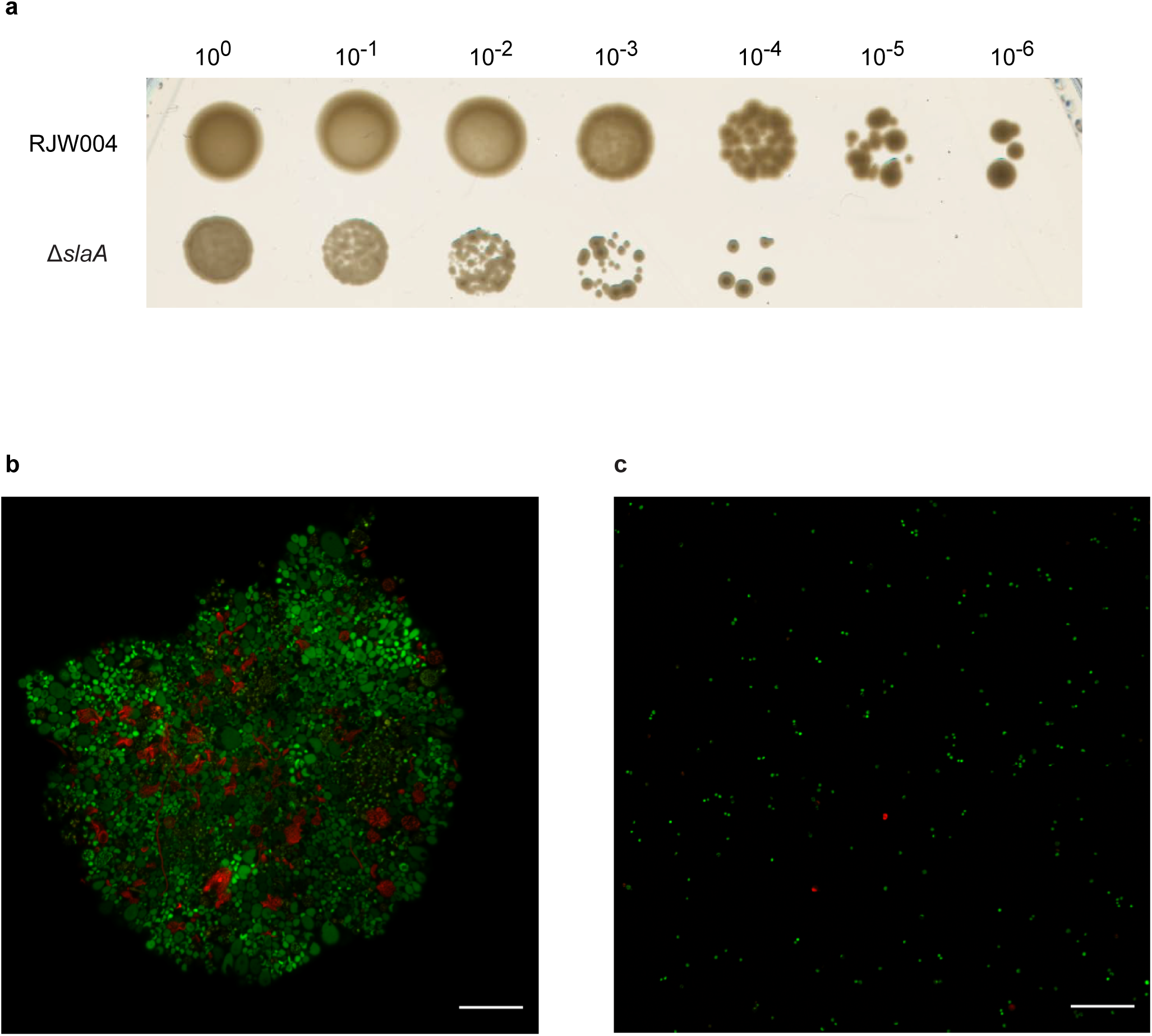
Cell viability of the wild-type RJW004 and ∆*slaA* mutant. A representative plate of the spotting assay **(a)**. Ten microliter of 10-fold serial dilutions of the wild-type RJW004 and ∆*slaA* mutant (OD_600nm_ was adjusted to 0.5) are spotted on the plate medium. The plates are incubated at 78 °C for 12 days, and then imaged with a scanner. Three biological replicates are performed for each sample in the spotting assay. Live-dead fluorescence microscopy of the ∆*slaA* mutant **(b)** and wild-type RJW004 **(c)**. Cells are stained with the Live/Dead BacLight Bacterial Viability Kit as described in “Materials and Methods”. Cells with intact and compromised membrane are labeled in green and red, respectively. Scale bars, 20 μm.

### *S. islandicus* cells deficient in S-layer exhibit a higher sensitivity to osmotic stress than the wild type but do not reduce in size

A theoretical analysis of the S-layer suggests that a crenarchaeal cell lacking an S-layer would have an increase in cell diameter when exposed to high internal pressure, such as turgor pressure (15). We hypothesize that, without the S-layer to constrain cell expansion and counteract the turgor pressure, the cells will become distended. To test this hypothesis, we grew wild-type RJW004 and ∆*slaA* mutant cells in liquid media supplemented with different concentrations of sucrose. This assay showed that wild-type strain was able to endure up to 2% (w/v) sucrose, in which the growth rate was still comparable to those of cells that grew in sucrose media with concentrations ranging from 0% to 1%; however, the wild type growth was severely impaired when cultivated in 5% sucrose media (Fig. 7a). In contrast, above 1% sucrose concentration, growth of the ∆*slaA* mutant was significantly delayed (Fig. 7b). These results showed that *S. islandicus* cells deficient in S-layer are more sensitive to higher osmotic stress when compared to wild-type cells. If large cells result from turgor pressure, expanding cells without an S-layer grown in media with higher concentrations of extra cellular solutes should relieve this pressure and result in smaller cell morphologies. Surprisingly, under higher osmotic conditions (1-5% of sucrose solution), aggregates containing large cells were still observed in ∆*slaA* mutant cells (Supplementary Fig. 9), indicating that cells are not expanding due to turgor pressure alone.

**Figure 7.**
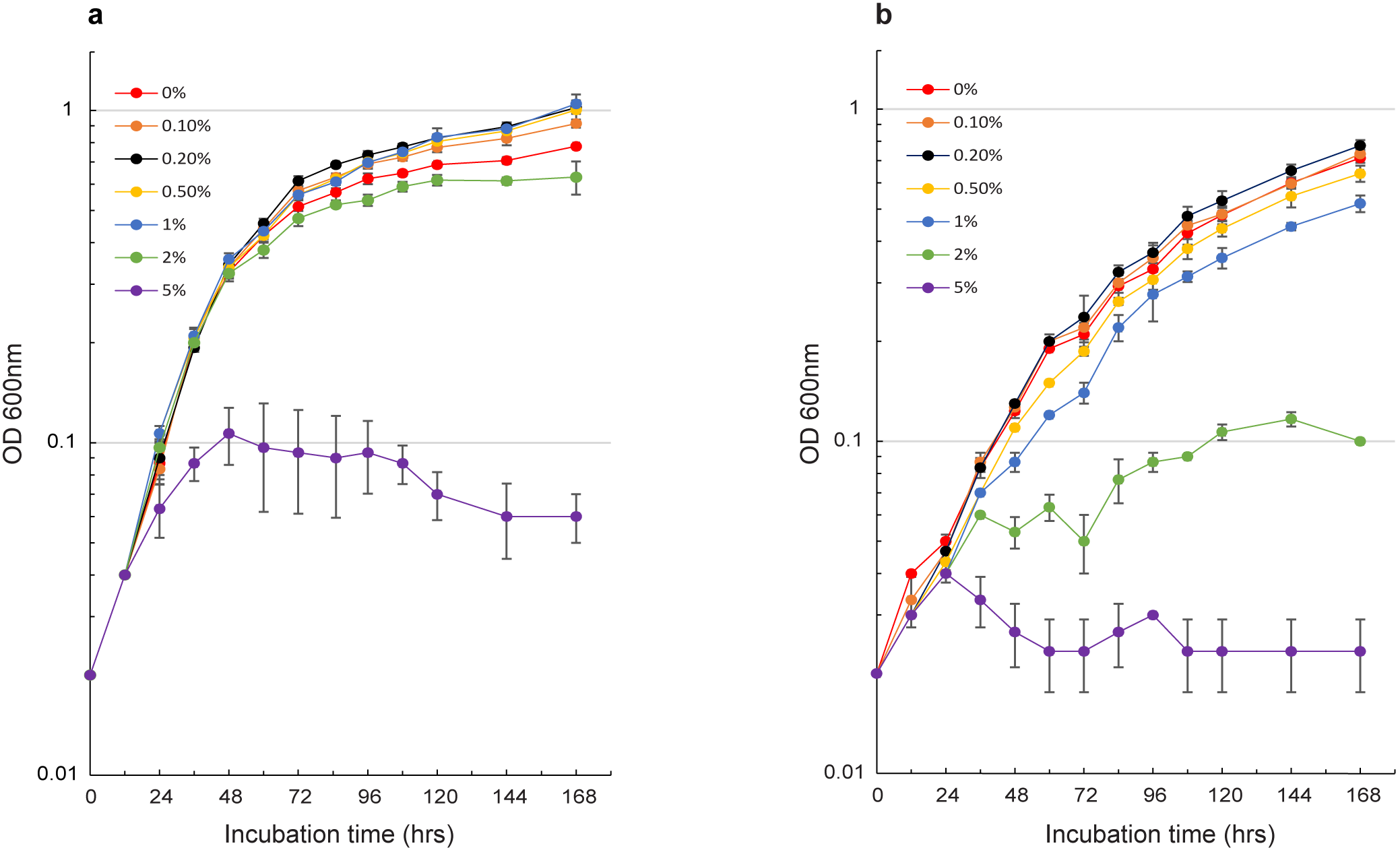
Growth profiles of wild-type RJW004 and ∆*slaA* mutant strains in dextrin-tryptone (DY) medium supplemented with different concentrations of sucrose. The wild-type strain RJW004 (**a**) and ∆*slaA* mutant (**b**) are cultivated at 78°C in the presence of 0%, 0.1%, 0.2%, 0.5%, 1%, 2%, and 5% sucrose solution, respectively. Cell growth is monitored for 7 days by measuring the OD_600nm_ of cell cultures. Error bars represent SD.

### *S. islandicus* cells deficient in S-layer exhibit variations in DNA content

*Sufolobus* cells with a cell division defect often have an enlarged cell size phenotype (32, 33). To investigate whether absence of the S-layer might impact cell division, wild-type RJW004 and ∆*slaA* mutant cells, taken from an exponential growth stage, were analyzed with flow cytometry. As shown in Fig. 8a, RJW004 exhibited a typical chromosomal pattern of the cell cycle, which was also observed in other *Sulfolobus* species (34, 35). However, in the ∆*slaA* mutant strain, the peaks in the fluorescence histogram corresponding to 3C (C refers to chromosome) and 4C as well as a long trailing tail became distinct, suggesting the existence of multiple chromosomes in the cells (Fig. 8b). It should be noted that the ∆*slaA* mutant cells tend to form big clumps in cultures, which interfere with interrogating the chromosome content of single cells using flow cytometry. Gating the single cells or including larger aggregates revealed that both have clear 3C- and 4C-peaks as well as a long tail with higher chromosome content (Supplementary Fig. 10). We note that in contrast to the wild-type cells, ∆*slaA* cells contain a greatly reduced population in the process of replicating between the 1C and 2C peak (Supplementary Fig. 10).

**Figure 8.**
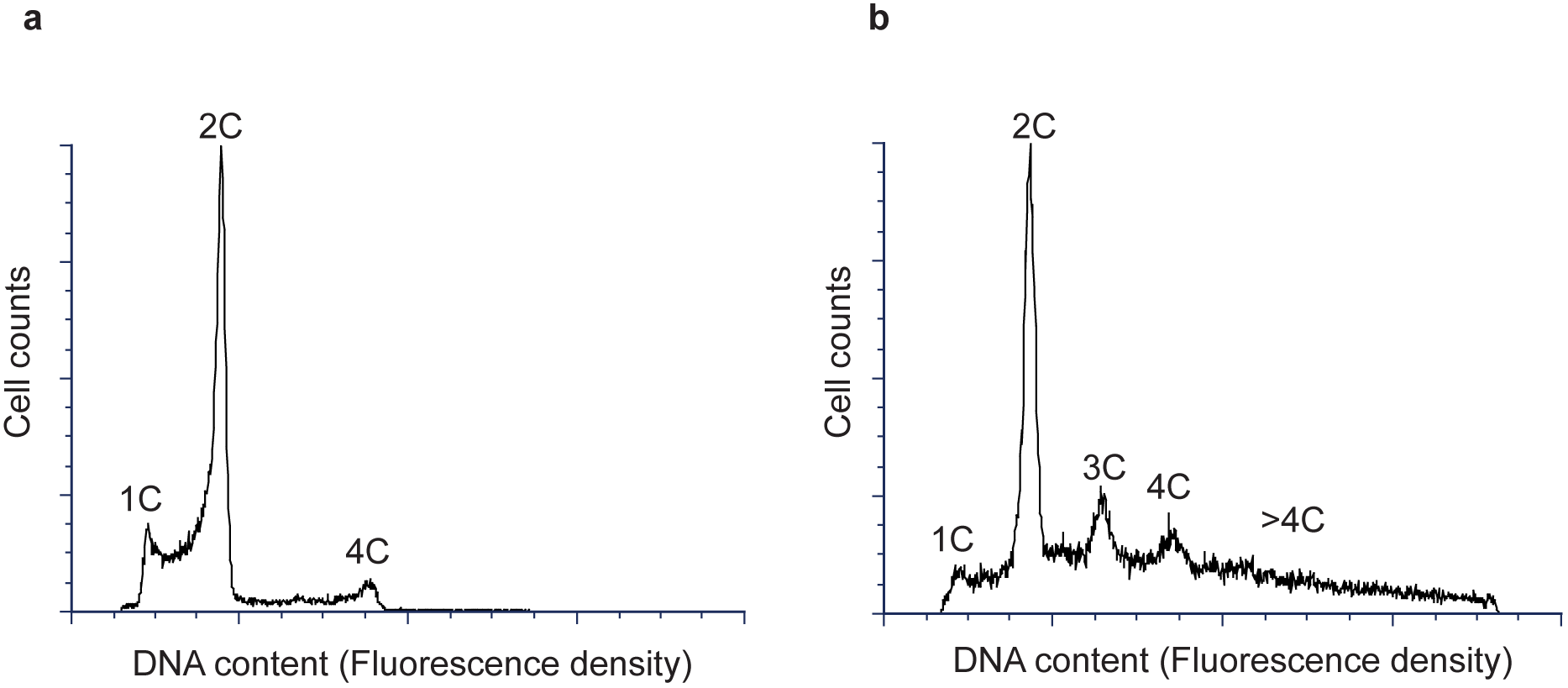
Flow cytometric profiles of the wild-type RJW004 (a) and ∆*slaA* mutant (b) strains. Cells from exponentially growing cultures were used. Cells with 1C, 2C, 3C, and 4C DNA content in the fluorescence histograms are indicated. C refers to chromosome.

## Discussion

The physiological and cellular functions of the archaeal S-layer remained largely unexplored, likely due to the experimental limitations or unfeasibility to create viable S-layer deficient mutants. Surprisingly, our recent work showed that archaeal cells lacking the S-layer are still able to maintain their cell viability in a crenarchaeon *S. islandicus* (10). In the present study, we have further characterized the S-layer gene deletion mutants, and provided strong genetic and cytological evidence to reveal that SlaA is the sole component that forms the outermost layer surrounding the whole cell whereas SlaB serves as a primary “stalk” to tether SlaA to the cell membrane. These results confirm our previous findings (10), and provide direct *in vivo* evidence in support of a so-called “stalk-and-cap” model of S-layer proposed in Sulfolobales (9, 36). Notably, we identified and genetically characterized a novel S-layer-related gene, *M164_1049*, encoding a SlaB paralog. The predicted transmembrane helix (Supplementary Fig. 3) present in both the N- and C-terminal of M164_1049 suggests that M164_1049 has a membrane topological organization, implying there is a potential connection between M164_1049 and the outermost layer (SlaA). Interestingly, the M164_1049 homolog in a related *Sulfolobus* species *S. solftaricus* P2 (SSO1175, sharing 83% of amino acid identify with 96% of query coverage) has been shown to be an N-glycosylated protein (37), and biochemically characterized to function as a multi-domain thermopsin-like protease (38). Our data show that M164_1049 is not required to maintain a structured and intact outermost layer (SlaA) when SlaB is present, as the single ∆*M164_1049* mutant exhibits a very similar S-layer structure as that of the wild-type strain. However, when SlaB was absent, SlaA became quite unstable, indicating that M164_1049, alone or even together with other S-layer-associated proteins, was not able to completely fulfill the roles of SlaB, the primary stalk. Moreover, in the absence of both SlaB and M164_1049 (∆*slaB*∆*M164_1049* mutant), the outermost layer was not seen outside of the cytoplasmic membrane, as observed in the electron micrographs. Collectively, these data show that M164_1049 serves as a secondary stalk and assists the primary stalk SlaB to anchor SlaA firmly to the cytoplasmic membrane. Given the broad distributions of M164_1049 within Sulfolobales and the role of M164_1049 in anchoring the SlaA lattice structure when SlaB is absent, though via a weak connection, it should be considered when rebuilding a new S-layer model in Sulfolobales. These data show that our understanding of interactions between membrane proteins and the crystalline layer SlaA are only just beginning to be explored.

*S. islandicus* cells that are deficient in SlaA, either caused by a genetic deletion of the *slaA* gene from the chromosome (∆*slaA* and ∆*slaAB* mutants) or by the physical absence of SlaA as in the ∆*slaB*∆*M164_1049* mutant cells, form large aggregates composed of cells with an increased diameter in liquid culture. The ∆*slaB* mutant also exhibits a high level of cellular aggregation (approximately 80% cells are in aggregates), but the sizes of aggregates along with the cell diameters are smaller than that of ∆*slaA* mutant. We proposed that the consequence of high level of cellular aggregation but with relatively small-size aggregates in the ∆*slaB* mutant is probably due to a large proportion of cells that retained a partial outermost layer (SlaA), as revealed by electron microscopy analysis of the ∆*slaB* mutant. We believe this retention of the partial SlaA in the ∆*slaB* mutants is also responsible for the slight increase in ∆*slaB* cell diameter compared to wild-type cells. These data suggest that SlaA, not SlaB, is the key determinant of cellular aggregation and cell morphology in *S. islandicus*, though the exact contribution of SlaB to these phenotypes remains to be investigated.

Remarkably, we do observe an inconsistency of cell-size range between the light microscopy and SEM for S-layer gene deletion mutants, with the cell size in the latter generally smaller. It is possible that the cells, without the protection of a functional outermost layer (SlaA), are not rigid and thus are flattened by the coverslip to some extent during light microscopy which possibly increased the observed cell diameter. Likewise, these SlaA^-^ cells may have suffered from serious damage during the SEM sample preparation, which may reduce cell size or cause the larger cells to not survive the treatment intact.

The molecular mechanisms that result in large cell clusters and cell size in SlaA^-^ are not clear. It has been shown that cellular aggregation in *Sulfolobus* species is mediated by elevated UV-induced pili (ups pili) formation, upon the treatment by UV irradiation or DNA damage reagents such as mitomycin C and NQO (39, 40). However, it is unlikely that the cellular aggregation formed in S-layer gene knockout mutants is mediated by increased pili formation because we did not observe abundant ups pili structures, which are usually very short and straight (39, 41), in any S-layer mutant during TEM experiments, suggesting other mechanisms are responsible for the aggregation phenotype. The large aggregates of cells could be forming due to some uncharacterized interactions between the now exposed outer cell membrane surfaces that are no longer protected by the protein shell SlaA; however, this idea needs to be further investigated. Additionally, our data suggests that the cells are not expanding due to turgor pressure alone because we continued to see large cells even under conditions where cells should not be experiencing high turgor pressure, such as high osmotic conditions. Therefore, there must be alternative explanations as to why the cells are expanding.

We suggest two hypotheses, a cell fusion and cell division defect model, to explain the combination of large cell sizes within aggregates and surprisingly variable odd numbers of chromosomes observed in the ∆*slaA* mutant. The first is a cell fusion model where the SlaA deficient cells merge together, resulting in an increase in cell size. As the cells continue to grow over time, a wide range of cell diameters would be created, as observed in the ∆*slaA* mutant. This idea is supported by the fact that despite the ∆*slaA* mutant and wild-type cultures having comparable growth rates (10), we observed a decrease in OD_600nm_ and colony forming units (cfus) for the ∆*slaA* mutant, which could be explained by the presence of the large aggregates. These large aggregates would result in more light passing through the liquid culture when measuring the optical density, resulting in a lower OD_600nm_ value, as observed for ∆*slaA*. An additional effect would be that these aggregated cells would grow as one colony forming unit (cfu) on solid media, consistent with the observation that there is a more than 10-fold reduction in the cfu count of the ∆*slaA* mutant in spotting assays. Combined with the light microscopy estimates of the number of cells within aggregates, this indicates that there is a high level of aggregation which would allow the cells to have prolonged and close contact with one another leading to fusion. However, we cannot exclude the possibility that the lower cfu count may be caused or contributed by other factors such as an increased sensitivity to vortexing and pipetting. In the cell fusion model the cell membranes of the “naked” cells within aggregates fuse, accounting for the increased chromosome copy number. We suspect that this type of hypothesis alone would result in mutants with a greater increase in even number chromosome copy numbers due to the higher chance of contact between the cells that are in an extensive G2 stage during the *Sulfolobus* cell cycle (35). However, this model does not well explain the cell populations with a distinct 3C-peak observed in flow cytometry analysis unless 1C cells are fusing with 2C cells before they replicate the chromosome.

Alternatively, cells without the S-layer could be deficient in cell division and/or cell cycle checkpoints in DNA replication. The larger cells observed in the ∆*slaA* mutant can be explained by a cell division defect as a similar phenotype has been observed in *Sulfolobus* cells with a disturbance or defect in cell division (32, 33). It is worth mentioning that the observed cell sizes of the ∆*slaA* mutant (1-9 μm) have a much larger range than those of cells with a cell division defect or disturbance reported so far. Though S-layer gene deletion mutants have been created and characterized in several bacteria (3-5, 42, 43), cell division defect phenotypes in those mutants have been rarely reported with few implications (3). This is probably due to the existence of a more important cell wall component i.e. the peptidoglycan in bacteria, which significantly decreased the importance of the S-layer. Although the representativeness of the flow cytometric profile for the ∆*slaA* mutant cells remains unclear, the existence of a significant cell population with more than two chromosomes suggests that *S. islandicus* cells deficient in the S-layer lost the ability to regulate the cell cycle. In the cell division defect scenario, the observation of cell populations with an obvious 3C-peak in the ∆*slaA* mutant can be explained by one daughter cell re-replicating while the other does not before the cell division occurs. This is very unusual for *Sulfolobus* cells where the cell cycle is tightly regulated (44). In addition, we observed a decreased number of cells transitioning between 1C and 2C (Supplementary Fig. 10), suggesting a possibility that cells were no longer dividing to 1C but replicating before the division was completed.

The cell fusion and cell division hypotheses are clearly not mutually exclusive, and both cell fusion and irregularities in the cell cycle may be linked to S-layer function. It should be noted that neither of these two hypotheses can well explain why cells aggregate in the SlaA^-^ strains. Addressing these questions will allow for a more complete understanding of the S-layer function in *Sulfolobus* and archaeal cell biology as a whole as well as interactions with fundamental processes such as chromosome partitioning.

## Materials and Methods

### Strains and growth conditions

The strains and plasmids used in this study are listed in Table 1. The *E. coli* Top10 (Invitrogen, USA) was used to maintain and propagate plasmid DNA, and grown in Luria-Bertani (LB) liquid medium at 37°C. When required, 100 μg/ml ampicillin was supplemented in LB agar plates to select ampicillin-resistant clones. *S. islandicus* RJW004, the genetic host used in this study, was originated from a wild-type isolate *S. islandicus* M.16.4 (45), which carried in-frame deletions of *pyrEF*, *lacS*, and *argD* (46). *S. islandicus* strains were cultured at 76-78 °C without shaking in Dextrin-Tryptone (DY) liquid medium (0.2%[w/v] dextrin; 0.1%[w/v] tryptone), as described previously(31). When necessary, 20 μg/ml uracil or/and 50 μg/ml agmatine were added. *S. islandicus* single colonies were isolated on Gelrite-solidified plates, formulated as described previously (46), via an over-lay cultivation approach (47). Plates were put into sealed plastic bags and cultivated for 10-20 days at 76-78°C. Cell growth was monitored by optical density measurements at 600 nm (OD_600nm_) with a CO8000 cell density meter (WPA, Cambridge, United Kingdom).

**Table 1.** Strains and plasmids used in this study.

| Strain or plasmid | Genotypes/Descriptions | Reference/Source |
| --- | --- | --- |
| **Strain** |  |  |
| *S. islandicus* RJW004 | $\Delta pyrEF \Delta lacS \Delta argD$ | (46) |
| *S. islandicus* RJW011 | $\Delta pyrEF \Delta lacS \Delta argD \Delta slaA$; refer to $\Delta slaA$ | (10) |
| *S. islandicus* RJW012 | $\Delta pyrEF \Delta lacS \Delta argD \Delta slaB$; refer to $\Delta slaB$ | (10) |
| *S. islandicus* RJW013 | $\Delta pyrEF \Delta lacS \Delta argD \Delta slaAB$; refer to $\Delta slaAB$ | (10) |
| *S. islandicus* RJW014 | $\Delta pyrEF \Delta lacS \Delta argD \Delta M164_{1049}::StoargD$; *M164_1049* was replaced by the *StoargD* marker cassette in RJW004; refer to $\Delta M164_{1049}$ | This study |
| *S. islandicus* RJW015 | $\Delta\text{pyrEF}\Delta\text{lacS}\Delta\text{argD}\Delta\text{slaB}\Delta\text{M164\_1049}::\text{StoargD}$;<br>*M164_1049* was replaced by the *StoargD* marker cassette in<br>RJW012; refer to $\Delta\text{slaB}\Delta\text{M164\_1049}$ | This study |
| **Plasmid** |  |  |
| pSeSd-StoargD | *Sulfolobus-E.coli* shutter vector pSeSd carrying the arginine<br>decarboxylase-expression cassette derived from *Sulfolobus*.<br>*tokodaii* | (31) |

### RT-PCR analysis

Ten milliliters of cell cultures taken from exponentially growing strains were harvested and lysed with 1 ml of TRIzol® Reagent (Invitrogen, USA). Total RNA was extracted and isolated with PureLink™ RNA Mini Kit (Thermo Fisher Scientific, USA), and then cleaned and concentrated with RNA Clean & Concentrator-25 Kit (Zymo Research, USA). To exclude potential DNA contamination, the total RNA (~2 μg) was treated with 2 μl of amplification-grade DNase I (1 U/μl; Invitrogen, USA). Approximately 500 ng of DNase I-treated total RNA was reverse transcribed using 2 μM of gene specific reverse primers and 1 U of SuperScript® IV Reverse Transcriptase (Thermo Fisher Scientific, USA) following the manufacturer’s guidelines. Two microliters of synthesized cDNA was used as a template in a standard PCR amplification reaction performed in a thermocycler (Bio-Rad, USA), using the Phusion® High-Fidelity DNA polymerase (NEB, USA) and the gene specific-primers binding the regions inside of *slaA* and *slaB* genes (Supplemental Table S2). Genomic DNA and total RNA were used as templates for positive and negative controls in PCR amplification, respectively. The resulting PCR products were separated by electrophoresis with a 1.2% (w/v) TAE agarose gel.

### Construction and verification of the *M164_1049* gene deletion mutants in *S. islandicus*

Deletion of *M164_1049* in the genetic background of RJW004 and RJW013 (Table 1) was accomplished by replacing it with an arginine decarboxylase expression cassette (*StoargD*) respectively, using a recently developed MMGI approach (31). Agmatine prototrophic (ArgD^+^) transformants were selected on DY plates containing 20 μg/ml of uracil but lacking agmatine. The *M164_1049* deletion mutants were verified by a colony PCR procedure described previously (46) with two primer sets: 1) the flanking primer set annealing outside region of the target gene, and 2) the internal primer set that specifically bound the inside region of the target gene. The resulting PCR products were treated with an ExoSAP-IT™ PCR Product Cleanup Reagent kit (Thermo Fisher Scientific, USA), and then sequenced to confirm the insertion of *StoargD* marker cassette in the desired position. *S. islandicus* mutant strains were purified at least once by an over-lay plating method (47), and then used for further studies.

### Phase contrast microscopy

*S. islandicus* strains were grown to the mid-log phase. Five microliters of cell cultures were spotted on a cleaned microscope slide using a blunted 20-μl pipette tip in order to avoid potential disruption of the large aggregates formed in the S-layer gene deletion mutants. A coverslip was added immediately, and then the slide was observed on a Zeiss Axiovert 200M microscope with a 63×/1.40 oil objective. Images were captured using a ZEISS Axiocam 506 microscope camera and visualized with Zeiss Zen imaging software. For quantitative cellular aggregation assays, cells were observed from at least 20 fields of the microscope slide and imaged to collect at least 1,000 single and, when possible, aggregated cells for each sample. Three biological replicates were set up for each sample.

### Electron microscopy

Samples used for conventional TEM analysis were prepared as follows. 200-mesh Carbon Type-B grids (TED Pella, INC.) were placed on 20-μl droplets of *S. islandicus* cell cultures for 3 minutes. The residual cell cultures on the edge of grids were adsorbed with a Whatman^TM^ filter paper, washed with degassed water, and then the grids were negatively stained with 2% uranyl acetate (w/v) for 15-30 seconds. Samples used for thin-section TEM analyses were essentially prepared as described by Bautista *et al* (48) with minor modifications (10). The grids were observed with a Philips CM200 transmission electron microscope operated at 120 kV. Images were taken using a Peltier-cooled Tietz (TVIPS) 2k × 2k CDD camera, and processed with EM-MENU software. For SEM analysis, 8-10 ml of cell cultures taken from the mid-log phase were filtered through a 0.2-μm filter (Whatman, USA) with a 10-ml syringe and then fixed at 4 °C for 4 hours in a fixative solution (2.0% paraformaldehyde and 2.5% glutaraldehyde in 0.1 M Sodium Cacodylate Buffer, pH 7.4). Afterwards, the fixation buffer was removed and replaced with a rinse buffer (0.1 M Sodium Cacodylate Buffer, pH 7.4). Samples were gently shaken for 10 min, and then gradually washed in 37%, 67%, 95%, and 100% (v/v) ethanol for 10 minutes each. Following the last step, the samples were washed twice more with the 100% ethanol, and the filters were critical point dried within 48 hours using the tourimis Critical Point Dryer (Autosamdri®-931). To observe the samples, filters were sputter coated with the Au-Pd and then imaged with a HITACHI S-4800 HIGH RESOLUTION SEM at 10 kV at various magnifications.

### Flow cytometry

Three hundred microliters of *S. islandicus* cells were fixed with 700 μl of ice-cold absolute ethanol, and the mixture was then briefly vortexed. Fixed samples were stored at 4 °C for at least 12 hours. Afterwards, the fixed cells were centrifuged at 13, 000 rpm for 5 min and resuspended in 1 ml of cold Tris-MgCl_2_ buffer (10 mM Tris-HCl, pH = 7.5, 10 mM MgCl_2_). Then the samples were precipitated and resuspended again in 300 μl of Tris-MgCl_2_ buffer containing 2 μg/ml of SYTOX Green nucleic acid stain (Thermo Fisher Scientific, USA) and 100 μg/ml of RNase A (DNase and protease-free, Thermo Fisher Scientific, USA). After at least 30 minutes’ incubation on ice in the dark, samples were analyzed using a LSR II (BD Biosciences, USA) flow cytometer, with a 488 nm (50 mW) blue laser as excitation light and a 530/30 band-pass filter. A total of 100,000 cell counts for each sample were collected. Flow cytometry data were processed and analyzed with De Novo FCS Express 6 software.

### Live-dead staining and microscopy

RJW004 and the ∆*slaA* mutant strain were grown to a mid-log phase. For the wild-type RJW004, 1 ml of cultures were then transferred to a 1.5-ml microcentrifuge tube and then centrifuged at 10, 000 rpm for 5 min to pellet. The pellet was resuspended with 1 ml 1×DY medium to wash. Centrifugation was repeated once, with the resulting cell pellet then resuspended in 500 μl of dye solution. For the ∆*slaA* mutant cells, to preserve the aggregate phenotype, a total of 5 microcentrifuge tubes containing 1 ml of cultures were allowed to settle for 15 min to collect cells at bottom of the tube. Cells of all 5 microcentrifuge tubes were then combined with supernatant removed. The remaining cells were resuspended in 1 ml of fresh DY liquid medium. Cells were allowed to settle for an additional 10 min. Supernatant was removed and the cells were gently resuspended in 500 μl of the dye solution, as per protocol (3 μl of 1:1 dye components A and B per 1000 μl 1×DY medium) (L7007 Live/Dead BacLight Bacterial Viability Kit, Invitrogen, USA). Cells were covered and incubated at room temperature in darkness for at least 15 min before viewing. Five microliters of stained-cells were then spotted directly on a microscope slide using a blunted 20-μl pipette tip, immediately covered with a coverslip, and observed on a Zeiss LSM 710 confocal microscope with 63×/1.40 Oil M27 objective at excitation wavelengths of 561 nm (for PI detection) and 488 nm (for SYTO 9 detection) using a ZEISS Zen software. A z-stack of the aggregates was taken at 1-μm slices.

### Osmotic pressure test

For the osmotic stress assay, 50% sucrose solution was added into the 45 ml of DY liquid medium, making the final concentrations (w/v) of sucrose with 0%, 0.05%, 0.10%, 0.2%, 0.5%, 1.0%, 2.0% and 5.0%. Proper volumes of *S. islandicus* cells were transferred into the medium to make an initial OD_600nm_ of 0.02. Three biological replicates were established for each treatment. Cells were grown at 78°C without shaking and their growth was monitored for 7 days by measuring the OD_600nm_ usually every 12 or 24 hours. Cell cultures taken from a designated OD_600nm_ were examined with a light microscope.

## Acknowledgments

We particularly thank Mohea Couturier and Elizabeth F. Rowland for insightful discussions and edits of the manuscript. Scott Robinson, Cate Wallace, and Charles Bee from Microscopy Suite at the Beckman Institute for Advanced Science and Technology, University of Illinois at Urbana-Champaign (UIUC), are greatly acknowledged for providing assistance with SEM and TEM sample preparation. We acknowledge Austin Cyphersmith from Core Facilities at the Carl R. Woese Institute for Genomic Biology, UIUC, for providing support with wide field and fluorescence microscopy imaging. We thank Barbara Krystyna Pilas and Barbara Maria Balhan from Flow Cytometry Facility at Roy J. Carver Biotechnology Center, UIUC, for help in choosing nucleic acid stains for flow cytometry, setting up flow cytometer parameters, as well as data interpretation.

## Author Contributions

C.Z., and R.J.W. conceived and designed the research; C.Z., R.L.W., Y.L., Z.W., and E.N.H. carried out experiments; C.Z., R.L.W., Y.L., and R.J.W. analyzed the data; R.J.W contributed new reagents/analytic tools and supervised the studies. C.Z., R.L.W., and R.J.W. wrote the paper. All authors edited the manuscript.

